# Antibiotic prophylaxis and hospitalization of horses subjected to median laparotomy: gut microbiota trajectories and abundance increase of *Escherichia*

**DOI:** 10.1101/2023.05.24.542119

**Authors:** Anne Kauter, Julian Brombach, Antina Lübke-Becker, Dania Kannapin, Corinna Bang, Sören Franzenburg, Sabita D. Stoeckle, Alexander Mellmann, Natalie Effelsberg, Robin Köck, Sebastian Guenther, Lothar H. Wieler, Heidrun Gehlen, Torsten Semmler, Silver A. Wolf, Birgit Walther

**Affiliations:** Advanced Light and Electron Microscopy (ZBS4), Robert Koch Institute, Berlin, Germany; Center for Infection Medicine, Institute of Microbiology and Epizootics, Freie Universität Berlin, Berlin, Germany; Veterinary Centre for Resistance Research (TZR), Freie Universität Berlin, Berlin, Germany; Equine Clinic, Surgery and Radiology, Freie Universität Berlin, Berlin, Germany; Institute of Clinical Molecular Biology, Christian-Albrechts-University of Kiel, Kiel, Germany; Institute of Hygiene, University of Münster, Münster, Germany; Institute of Hygiene, DRK Kliniken Berlin, Berlin, Germany; Pharmaceutical Biology, Institute of Pharmacy, Universität Greifswald, Greifswald, Germany; Robert Koch Institute, Berlin, Germany; Genome Sequencing and Genomic Epidemiology (MF2), Robert Koch Institute, Berlin, Germany; Section Microbiological Risks (1.4), Department II Environmental Hygiene, German Environment Agency, Berlin, Germany

**Keywords:** Horse, microbiome, gastrointestinal tract, microbiota, 16S rRNA gene sequencing, hospitalization, colic, *Escherichia*

## Abstract

Horse clinics are hotspots for the accumulation and spread of clinically relevant and zoonotic multidrug-resistant bacteria, including extended-spectrum β-lactamase producing (ESBL) Enterobacterales. Although median laparotomy in cases of acute equine colic is a frequently performed surgical intervention, knowledge about the effects of peri-operative antibiotic prophylaxis (PAP) based on a combination of penicillin and gentamicin on the gut microbiota is limited. Therefore, we collected fecal samples of horses from a non-hospitalized control group (CG) and from horses receiving either a pre-surgical single-shot (SSG) or a peri-operative 5-day (5DG) course of PAP. To assess differences between the two PAP regimens and the CG, all samples obtained at hospital admission (t_0_), on days three (t_1_) and ten (t_2_) after surgery, were screened for ESBL-producing Enterobacterales and subjected to 16S rRNA V1– V2 gene sequencing.

We included 48 samples in the SSG (n=16 horses), 45 in the 5DG (n=15) and 20 in the CG (n=10). Two samples (6.5%) were positive for ESBL-producing Enterobacterales at t_0_ while this rate increased to 67% at t_1_ and decreased only slightly at t_2_ (61%). Shannon diversity index (SDI) was used to evaluate alpha-diversity changes, revealing that horses suffering from acute colic seemed to have a compromised fecal microbiota composition (5DG, SDI_mean_ of 5.90; SSG, SDI_mean_ of 6.17) when compared to the CG (SDI_mean_ of 6.53) at t_0_, although the difference lacked significance. Alpha-diversity decreased significantly in both PAP groups at t_1_, while at t_2_ the onset of microbiome recovery was noticed. Although we did not identify a significant SDI_mean_ difference with respect to PAP duration, the community structure (beta-diversity) was considerably restricted in samples of the 5DG at t_1_, most likely due to the ongoing administration of antibiotics. An increased abundance of *Enterobacteriaceae,* especially *Escherichia*, was noted for both study groups at t_1_. Further studies are needed to reveal important factors promoting the increase and residency of ESBL-producing Enterobacterales among hospitalized horses.

## 1 Introduction

Compared with other companion animals, horses more often acquire gastro-intestinal tract (GIT) disorders that may lead to long-term suffering or even death (Traub-Dargatz et al., 2001). The syndrome complex pain caused by disorders of the GIT in horses is commonly referred to as “colic” (Traub-Dargatz et al., 2001;Stockle et al., 2018). The composition of the bacterial community residing within the GIT has been regarded as beneficial and a prerequisite for the health and well-being of hindgut fermenters such as Equidaes (reviewed in (Kauter et al., 2019)). Previous reports indicated that colonization with the physiological endogenous microbiota shields the equine GIT against either direct or indirect pathogen-induced damages and that these protective effects are perturbed throughout various enteral maladies (Costa et al., 2012;Weese et al., 2015). Moreover, administration of antibiotics such as penicillin (Baverud et al., 2003), enrofloxacin or ceftiofur (Liepman et al., 2022), as well as doxycycline (Davis et al., 2006) were reported to drive the enteral microbial community towards a dysbiotic state (Costa et al., 2015). However, besides other factors, the state of the microbiota at the time of (antibiotic) perturbation (diet, species, and functional diversity and redundancy) and the characteristics of the perturbation (e.g. administration route, antimicrobial spectrum, and duration of antibiotic courses) determine the extent of any eventual dysbiosis (Schwartz et al., 2020).

To prevent adverse postoperative events such as surgical site infections in horses subjected to abdominal surgery due to acute colic, peri-operative antibiotic prophylaxis (PAP) is recommended (Dallap Schaer et al., 2012). The most commonly used PAP regimen for horses requiring median laparotomy consists of a combination of penicillin and gentamicin (P/G) for a period of 3-10 days (Dallap Schaer et al., 2012;Wormstrand et al., 2014;Teschner et al., 2015). In contrast, the standard regimen for similar surgical interventions lacking complicating circumstances in human and small animal medicine is a short-term PAP, which is provided as a single-shot therapy 30-60 minutes prior to incision (Stöckle et al., 2021). To investigate the effects of prolonged administration beyond this immediate peri-operative timeframe (> 24 h after surgery), we conducted a pilot study focusing on the clinical outcome (Stöckle et al., 2021), the extended-spectrum β-lactamase (ESBL)-producing *Escherichia coli* (EC) carriage rates (Kauter et al., 2021) and the microbiome composition (present study). Beyond others, our results indicated non-inferiority of a “single-shot” versus a five-day course of P/G PAP with respect to the patients’ clinical outcomes (Stöckle et al., 2021). Regardless of the applied PAP-regimen, we noticed a worrisome increase of ESBL-EC carriage rates among these horses during their hospital stay (Kauter et al., 2021).

Based on these previous observations, the current study aimed i) to examine the extent of microbiome disturbance in hospitalized horses subjected to median laparotomy, ii) to reveal alterations of the gut microbiota caused by a short and a long-term PAP regime and iii) to enable insights into changes of the microbiome that might play a role in the previously reported increased ESBL-EC carriage rates among equine patients.

## 2 Materials and Methods

### Study cohort and perioperative antibiotic prophylaxis

Horses diagnosed with acute abdominal pain (colic syndrome complex) that required median laparotomy were included in this study (detailed description of the controlled and randomized pilot study in (Stöckle et al., 2021)). Briefly, horses allotted to the SSG received short term P/G PAP, while the 5DG group received P/G PAP for five consecutive days. In both groups, PAP consisted of parenteral administration of sodium penicillin G (22,000 IU/kg four times daily) and gentamicin (6.6 mg/kg), as previously recommended for colic surgery (Dallap Schaer et al., 2012;Durward-Akhurst et al., 2013;Southwood, 2014). Specimens of ten non-hospitalized farm horses that were free of clinical symptoms for any apparent illnesses served as a control group (CG). The latter were sampled twice within three days to ensure representativeness of the results (detailed description of the study participants and their respective clinical outcomes in (Stöckle et al., 2021)). A graphical abstract of the study outline is provided in **Figure 1**.

**Figure 1.**
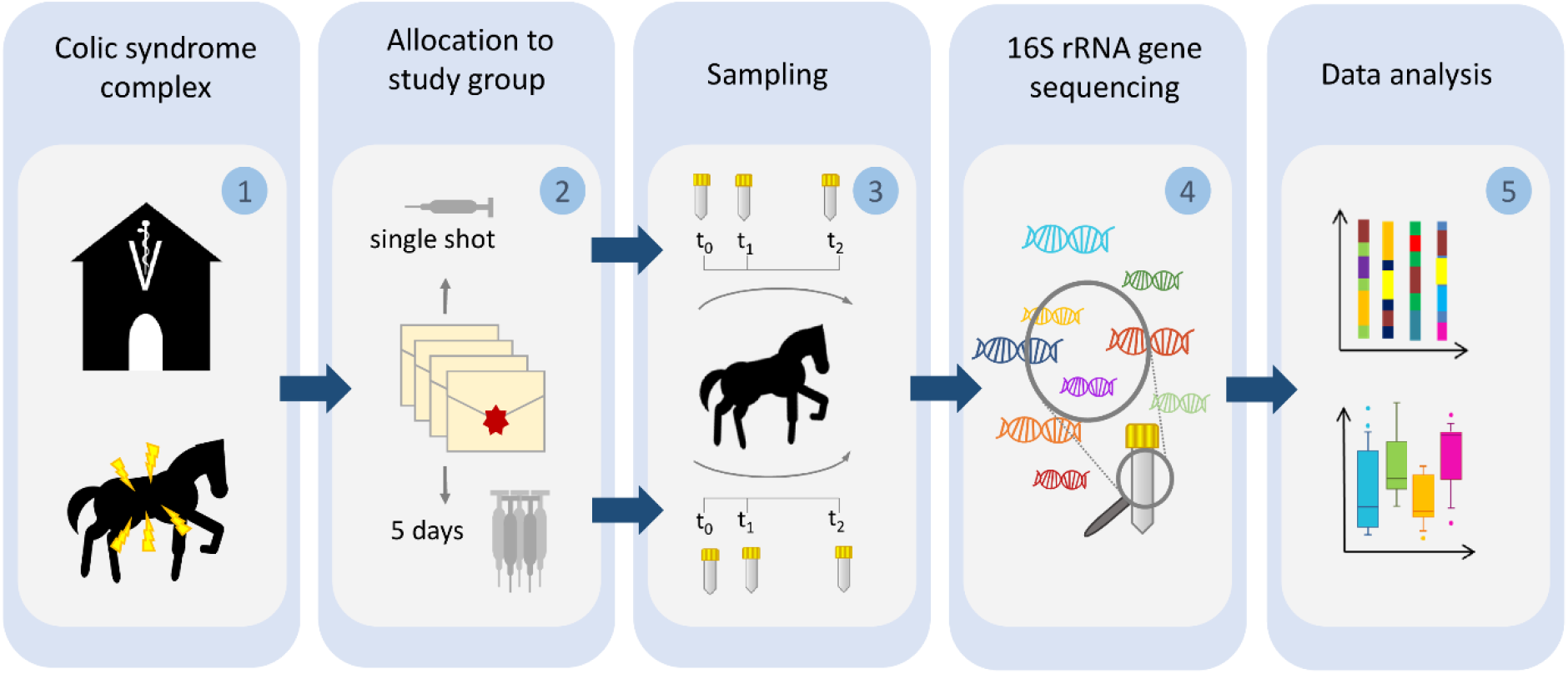
Illustration of the sampling procedure. Horses subjected to colic surgery were **(1)** sampled directly at hospital admission (t_0_) and **(2)** allotted to either a single-shot perioperative antibiotic prophylaxis regimen or a 5-day-lasting protocol. Sampling was repeated for all horses on day 3 (t_1_) and 10 (t_2_) after surgery **(3)**. All samples were subjected to DNA extraction and V1–V2 16S rRNA gene sequencing **(4)**. Sequences were preprocessed and analyzed with respect to changes of microbiota composition, alpha and beta diversity **(5)**.

Inclusion criteria for study participation were: study participants had to be free of clinical signs of infectious diseases prior to surgery. Additionally, since the juvenile microbiome is known for continuing changes during the foals’ gut maturation (Costa et al., 2016), all included horses were required to be one year or older. Equine patients were excluded from further consideration if their hospital stay had ended prematurely due to euthanasia and/or the antibiotic regimen they had originally been assigned to was not strictly followed, regardless of the particular reasons requiring these changes. Horses with incomplete sample sets (t_0_, t_1_, t_2_, see sample collection) were also excluded (Stöckle et al., 2021).

### Sample collection

Fresh fecal samples were collected from each horse diagnosed with colic syndrome complex directly at hospital admission (t_0_), as described previously (Walther et al., 2018;Kauter et al., 2021). A second sample was collected after three days (t_1_) and a third (final) sample was obtained ten days after surgery (t_2_). In order to gain insights into the gut microbiome associated with non-hospitalized horses lacking clinical signs of gastro-intestinal disorders, control samples were obtained from ten horses residing in a barn. All specimens were stored directly at -80° C and shipped on dry ice.

Identification of ESBL-producing Enterobacterales was previously described (Kauter et al., 2021). Briefly, samples were cultured on Brilliance™ ESBL Agar plates (Thermo ScientificTM, Germany) overnight. Colonies showing characteristic growth signatures of ESBL-producing Enterobacterales on chromogenic screening plates were further investigated. In case of distinct phenotypic appearances of presumptive ESBL-producing colonies on the plates, all isolates were subjected to an ESBL confirmation test according to the Clinical Laboratory Standards Institute’s (CLSI) recommendations (CLSI, 2020). Species confirmation was achieved by matrix-assisted laser desorption/ionization-time of flight (MALDI-TOF) mass spectrometry (Bruker, Germany).

### DNA extraction and sequencing of the bacterial 16S rRNA V1-V2 region

The sequencing of bacterial DNA was performed by the Institute of Clinical Molecular Biology (IKMB) at the Christian-Albrechts University of Kiel. DNA was extracted using the QIAamp Fast DNA stool mini kit (QIAGEN, Hilden, Germany) automated on the QIAcube (QIAGEN, Hilden, Germany). For this, approximately 200 mg feces were transferred to 0.70 mm Garnet Bead tubes filled with 1 ml InhibitEx buffer. Subsequently, bead beating was performed using a SpeedMill PLUS (QIAGEN, Hilden, Germany) for 45 s at 60 Hz. Samples were then heated to 95° C for 5 min and afterwards centrifuged for 1 min at 10,000 rpm. 200 µl of the resulting supernatant were transferred to a 1.5 ml microcentrifuge tube, which was placed in the QIAcube for follow-up automated DNA isolation according to the manufacturer’s protocol. DNA bound specifically to the QIAamp silica-gel membrane while contaminants passed through. PCR inhibitors were removed through means of InhibitEX (QIAGEN, Hilden, Germany), a unique adsorption resin, and an optimized buffer. DNA was diluted 1:10 prior to PCR, and 3 µl were used for further amplification. PCR-products were verified using agarose gel electrophoresis. PCR-products were then normalized using the SequalPrep Normalization Plate Kit (Thermo Fischer Scientific, Waltham, MA, USA), pooled equimolarily and sequenced on an Illumina MiSeq v3 2×300 base pair (bp) platform (Illumina Inc., San Diego, CA, USA).

The V1-V2 region of the 16S rRNA gene was subsequently sequenced (Primerpair 27F-338R, dual MIDs inducing) on a MiSeq-platform (MiSeq Reagent Kit v2) (Kozich et al., 2013). The resulting MiSeq raw fastq data was verified using an inhouse pipeline (Bcl2fastq Modul in CASAVA 1.8.2, Demultiplexing, FLASH software (v1.2) (Magoč and Salzberg, 2011), fastx toolkit und UCHIME (v6.0) (Edgar et al., 2011)). Demultiplexing was performed based on 0 mismatches within the barcode sequences.

### Analysis of 16S rRNA gene sequences

#### Data Preprocessing

Sequence reads were preprocessed as described (Mach et al., 2020). In brief, paired-end reads were merged into continuous sequences using the “join_paired_ends.py” script of QIIME (v1.9.1) (Caporaso et al., 2010) in “fastq-join” mode (Aronesty, 2013). Defined parameters allowed a minimum overlap of 6 bp and a maximum difference within the overlap region of 8%. Reads which did not meet these criteria were removed from further analysis. Next, seqkit (v0.16.1) (Shen et al., 2016) was utilized to filter out reads which were too short (≤300bp) or too long (≥470bp). The remaining sequences were then quality filtered using the “split_libraries_fastq.py” script of QIIME (v1.9.1) [25] by applying a PHRED quality threshold of 20. Reads were hereby required to have 50% of their bases to be consecutively of high quality. A maximum of three consecutive low-quality bases were allowed before truncating a read. Reads containing any ambiguous bases (“N”) were removed from further analysis.

#### OTU Clustering

Reads were clustered into operational taxonomic units (OTUs) using USEARCH (v11.0.667) (Edgar, 2010) and the Greengenes database (release 2013-08, gg_13_8_otus, 99_otus) (DeSantis et al., 2006) with a 97% similarity cutoff. The “unnoise3” algorithm of USEARCH was utilized for additional filtering of chimeric OTUs. In order to improve taxonomic annotation, Greengenes OTUs as well as unmatched representative sequences were then mapped against the RDP database (v11.5) (Cole et al., 2014) using the SequenceMatch pipeline of rdptools (v2.0.3) (Cole et al., 2014), searching for the closest neighbor (k=1) with a minimum sab score of 0.5. Counts were subsequently merged and the “filter_otus_from_otu_table.py” script of QIIME (v1.9.1) (Caporaso et al., 2010) was utilized to remove any singletons from the table (counts ≤ 3). Data were then exported using the biom (v 2.1.7) (McDonald et al., 2012) package and converted into the appropriate format for further data analysis. The resulting OTU table was then finalized by updating the available taxonomic labels with the recently released, new taxonomic names for bacterial phyla (Oren and Garrity, 2021).

#### Sample size calculation

Sample-size calculation was performed as recommended for microbiome studies by utilizing a permutation-based extension of the multivariate analysis of variance (PERMANOVA) on a matrix of pairwise distances (Kelly et al., 2015). Subsequent study power estimation was performed through the R package micropower (v0.4) (Kelly et al., 2015). Unfiltered OTU tables were hereby subjected to random rarefaction in order to assess key population parameters, including mean and standard deviation per OTU based on both the presence/absence of individual taxa as well as their abundance (Weighted Jaccard Distance) for a comparison of two groups of fixed size (n=10). These parameters were then further utilized to simulate a range of distance matrices and effect sizes in order to estimate the statistical power for identifying an effect size given a specified sample size and respective OTU table. Permutation-based sample-size estimation, according to Kelly et al. (Kelly et al., 2015), revealed that a subset of samples from ten horses per study group were sufficient to identify differences in taxonomic composition of effect size 0.020 with p=0.05 and a power of 80% (90% to identify an effect size of 0.035).

#### Diversity estimation and OTUs assignation

The resulting OTU counts were randomly sub-sampled for each sample to a homogeneous level, defined by the counts of the lowest sample (12,178). OTU counts above 10,000 have been shown to provide adequate comparisons between differing sequencing depths for microbiome analyses (Mach et al., 2020). Rarefaction was performed by using the “rarefy_even_depth” function of phyloseq (v1.36.0) (McMurdie and Holmes, 2013) with rngseed=1 in R (v4.1.1). Rarefied data was then utilized to assess the influence of antibiosis on the microbiome diversity among the equine patients and to visualize the distribution of taxa (OTUs) across the sample set. Within-sample diversity (alpha-diversity) was assessed through Shannon diversity indices (SDI) calculated using the R package microbiome (v1.14.0) (Lahti et al., 2017)). Between-sample diversity (beta-diversity) was determined through the computation of Bray-Curtis distances by phyloseq (v1.36.0) (McMurdie and Holmes, 2013) on all OTUs of the rarefied table. The resulting diversity metrics were further visualized with ComplexHeatmap (v2.8.0) (Gu et al., 2016) and ggplot2 (v3.3.5) (Wickham et al., 2016) for subsequent comparisons between the study groups. Non-parametric Paired Wilcoxon Rank Sum tests were performed between groups of interest and results with a p<0.05 were labeled as being significant. The Benjamini-Hochberg procedure was utilized for multiple-testing correction in order to limit the false discovery rate were applicable. OTUs were aggregated at selected taxonomic levels (including phlya) using the “aggregate_top_taxa” function of microbiome (v1.14.0) (Lahti et al., 2017). Differential taxa were identified between the PAP study groups (SSG / 5DG) using the raw OTU table and the microbiomeExplorer R package (Reeder et al., 2021) with proportional normalization and the DESeq2 method (Love et al., 2014). Additional correlation analyses were also performed within the microbiomeExplorer suite.

## 3 Results

To explore the effects of both G/P PAP regimens and hospitalization on the gut microbiota of horses subjected to median laparotomy, microbiome sequencing and analysis were performed using sample sets obtained from a pilot study comparing the clinical outcomes of 67 patients that met the study’s inclusion criteria (Stöckle et al., 2021). Overall, 48 samples from 16 horses representing the SSG and 45 samples of 15 horses belonging to the 5DG were comparatively evaluated. As a non-hospitalized CG we included additional 20 samples of ten horses that lacked any apparent signs of clinical illness, resulting in 113 samples (**Supplementary Table 1)**. At hospital admission, two samples (6.5%) were positive for ESBL-producing Enterobacterales (**Supplementary Table 1**), while, regardless of the study group, the overall rate increased to 67% (t_1_) and, only slightly decreased, at t_2_ (61%). There was no difference in carriage of ESBL-producing Enterobacterales between the SSG and 5DG (Fisher’s Exact Test, p = 1), while all CG samples were negative (Fisher’s Exact Test, p < 0.0001).

In total, 4,896,645 high quality OTU counts (ranging between 12,178 – 220,454 counts per sample, median = 32,730) were obtained and assigned to 17,035 different OTUs across 330 different taxonomic entities at genus level. Further taxonomic assignment of these OTUs revealed the top ten bacterial taxa (phylum level) based on their total counts across the sample set: The phyla *Bacteroidota* (38%) (previously *Bacteroidetes*) (Oren and Garrity, 2021) and *Bacillota* (33%) (previously: *Firmicutes*) (Oren and Garrity, 2021) were predominant in the equine fecal samples at hospital admission (t_0_), followed by *Verrucomicrobiota* (11%), *Pseudomonadota* (9%) (previously *Proteobacteria*) (Oren and Garrity, 2021) and *Spirochaetota* (4%) (**Figure 2**).

**Figure 2.**
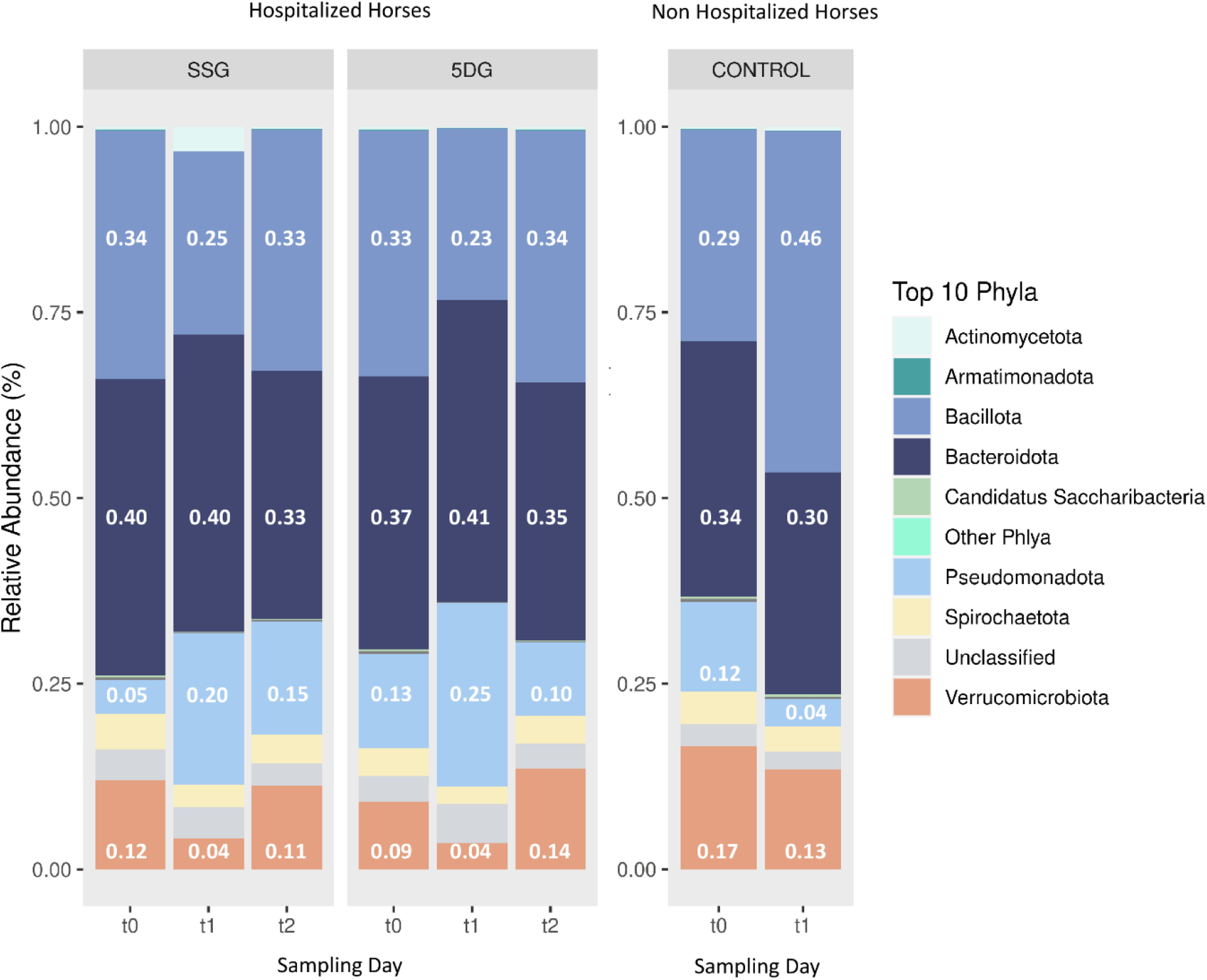
Taxonomic composition of microbiota in hospitalized horses. Stacked bar charts illustrating the relative abundances of the top ten phyla for the single-shot group (SSG, n=16) and the 5-day group (5DG, n=15). Abbreviations: t_0_ = hospital admission (SSG/5DG) / first sampling round (CG); t_1_ = three days after surgery (SSG/5DG) / three days after first sampling (CG); t_2_ = ten days after surgery (SSG/5DG).

Each horse harboured an individual composition of fecal bacterial communities, as presented in **Supplemental Figure 1**.

### i) Evaluation of gut microbiota profiles, microbiome disturbance, biodiversity and microbiota trajectories

#### Microbiota profiles at hospital admission

The SDI is a measure of a sample’s diversity based on both, community richness and -evenness. In the present study, SDI was selected to inspect the alpha-diversity within each sample and to compare these across study groups. Overall comparableness of the treatment groups was ensured by demonstrating the lack of significant difference in the mean SDI between both study groups at t_0_ (Wilcoxon test, P > 0.05). As expected, visualization of beta-diversity (“between sample diversity”) using ordination plots generated through principle component analysis (PCA) demonstrated an increased range of variance regarding the gut microbiota profiles of individual horses belonging to the SSG and the 5DG compared to the overall variance noticeable between the data points representing the CG samples (**Figure 3A-C**).

**Figure 3.**
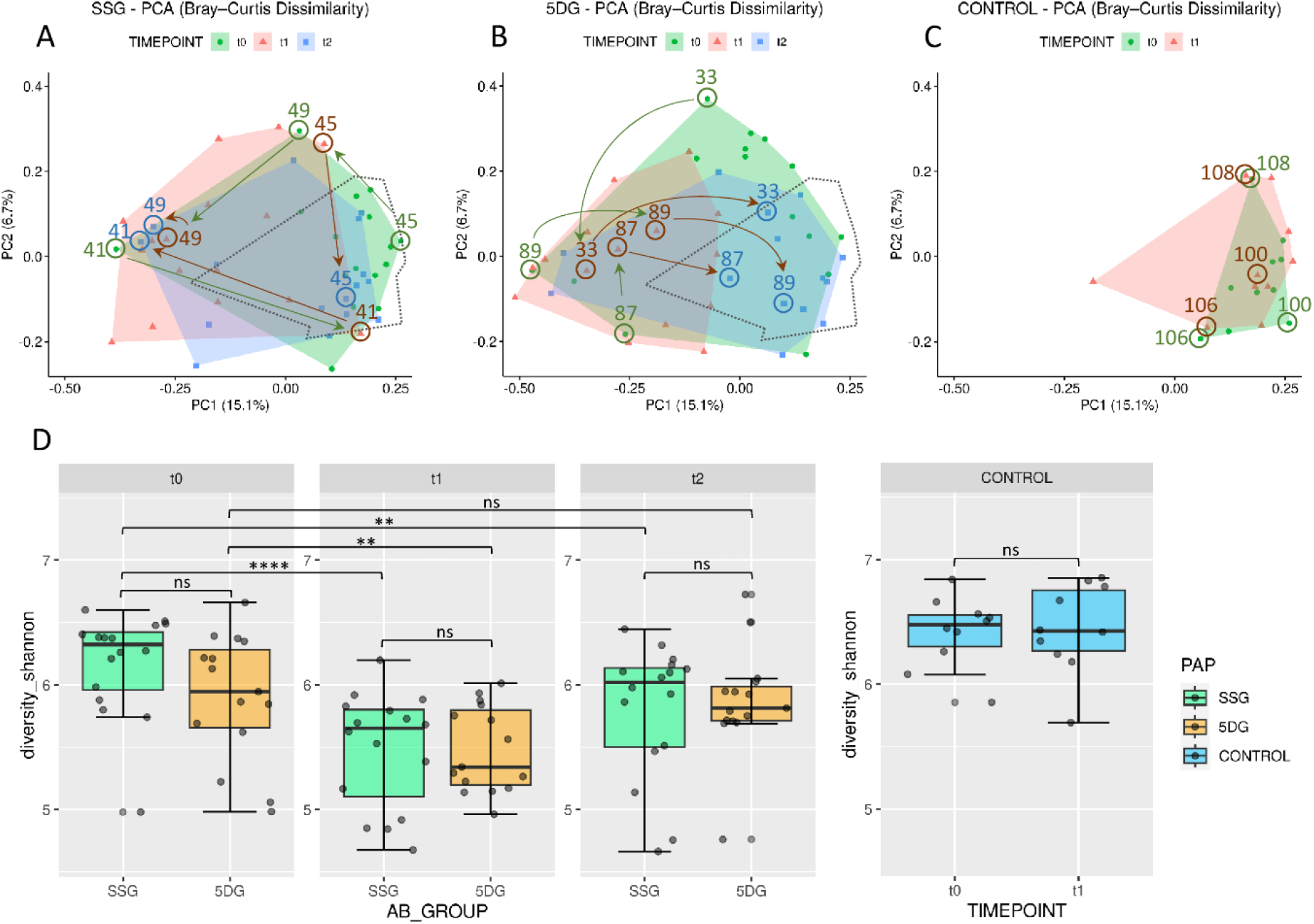
Diversity of the equine gut microbiome of hospitalized and non-hospitalized horses. **A-C)** PCA plots illustrating differences in gut microbiome composition (beta-diversity) based on unweighted Bray-Curtis dissimilarities across each group per sampling day. Axes represent the two dimensions that explain the largest proportion of variation across communities of each analysis. Distances between two data points reflect their similarity/dissimilarity with respect to sample composition. The colored areas represent the computed convex hull of data points from each sampling day, illustrating respective areas of minimal size. The area encompassing data points from the control group (CG) samples **(C)** is marked by a dotted line in **(A)** and **(B)** for direct comparison. Horses treated with peri-operative antibiotic prophylaxis (PAP) in the single-shot group (SSG) vs. the 5-day course group (5DG) show both large, intraindividual differences at t_0_ **(A)** and a strong shift away from the area defined by the CG samples at t_1_ **(B)**. Samples of both study groups converge back to the CG area at t2. Numbered arrows indicate the trajectories of microbiota composition within selected horses. **(D)** Boxplots displaying the alpha-diversity indices (Shannon) for all three groups (SSG, n=16; 5DG, n=15; control, n=10) (significance between groups: unpaired Wilcoxon Rank Sum tests, ** p<0.05, **** p<0.0005, ns=not significant).

Of note, the mean SDI values of the CG samples were 6.42 and 6.44, respectively. At t_0_, most of the enteral microbiota profiles associated with SSG horses (11 of 16) clustered within the framework set by the CG samples (**Figure 3A**). The t_0_ microbiota profiles obtained for the 5DG, on the other hand, illustrated increased variance among the 5DG samples, and only four out of 15 samples clustered with the CG area, potentially indicating that more horses belonging to the 5DG had, in comparison to the SSG, gut microbiota perturbances at hospital admission (**Figure 3B**). This observation is supported by our examination of the alpha-diversities, since the t_0_ samples representing the SSG were associated with an SDI_mean_ of 6.17, while the 5DG yielded an SDI_mean_ of 5.90, although the difference lacked statistical significance (**Figure 3D**).

#### Microbiota profiles at t_1_

At t_1_, most of the microbiota profiles obtained from fecal samples of the SSG (horses without antibiotics for at least 36h) and the 5DG (horses at the third day of their 5-day course P/G PAP) visibly differed from those obtained at t_0_ by shifting away from the centre of the CG area, indicating perturbations (**Figure 3B**). This shift was accompanied by a significant reduction in mean alpha-diversity across both groups [SSG: SDI from t_0_ = 6.17 to t_1_ = 5.48 (paired Wilcoxon Test, p=0.000031); 5DG: SDI from t_0_ = 5.90 to t_1_ = 5.48 (paired Wilcoxon Test, p=0.012)] (**Supplemental Table 1**, **Figure 3D**). In addition, the overall distances between 5DG samples seemed considerably restricted compared to the situation at t_0_ (**Figure 3B**). Samples representing the SSG, on the other hand, were more widely scattered than samples representing the 5DG or even the CG, indicating an increased level of inter-individual differences among microbiome compositions for SSG horses at t_1_.

#### Microbiota profiles at t_2_

At t_2_, i.e. ten (SSG) and five days (5DG) after the final PAP course was administered, the mean alpha-diversity of both study groups increased (SSG, SDI_mean_ = 5.80; 5DG, SDI_mean_ = 5.87), indicating the onset of microbiome recovery (**Supplemental Table 1**, **Figure 3C)**.

#### Common trajectories and inter-individual differences in microbiota profiles

To likewise illustrate common spatial shifts and individual deviation from common trajectories, data points representing samples of three individuals per group are highlighted in **Figure 3**:

**5DG**: At t_0_, the data points belonging to samples of equine patients 89, 33 and 87 clustered most distantly from the CG samples, indicating a considerable disturbance of the microbiome structure and composition at hospital admission. This finding is supported by the individual samples’ low SDI values (5.05, 5.75 and 5.29) (**Supplemental Table 1**). At t_1_, samples 33 and 87 showed a different composition compared to the t_0_ situation, but both points still clustered distantly from the CG area. Only the t_1_ sample obtained from horse 89 showed some signs of movement towards the CG area (**Figure 3B**). At t_2_, data points representing the samples of horses 33, 87 and 89 clustered within the CG area, indicating the onset of microbiome recovery that was accompanied by a notable SDI increase (5.76, 6.02, 5.76).

**SSG**: The initial data point representing the t_0_ sample of horse 45 (SDI 6.66) clustered near the area covered by the CG. While the corresponding data point at t_1_ indicated an increase of gut microbiome disturbance accompanied by an SDI decrease to 5.97, the t_2_ sample (SDI 6.15) clustered within the area framed by data points of the CG samples, once again indicating the onset of microbiome recovery.

However, some equine patients deviated from the aforementioned common temporal trajectory: The data points of horse 41 (SSG), for instance, indicated a considerable microbiome disturbance at t_0_ (SDI 5.05) followed by a brief relative recovery at t_1_ (SDI 5.99). Then, the data point of horse 41 shifted towards the opposite direction at t_2_ (**Figure 3A**), a reversal that is accompanied by an SDI decrease to 5.59.

Taken together, the control samples seemed to represent an overall beneficial structure and composition of equine gut microbiota, since all patients’ samples associated with an SDI > 5.8 clustered near or within the area covered by these samples, while samples associated with lower SDIs clustered elsewhere (**Figure 3** and **Supplemental Table 1**). Moreover, although a common temporal trajectory from hospital admission towards discharge was notable for the gut microbiota of most participants in both study groups, the temporal patterns of some horses deviated from these, emphasizing the individual nature of the GIT microbiome recovery process.

### ii) Evaluation of gut microbiota alterations among horses receiving different PAP regimens after colic surgery

To gain insights into the putative effect of the different PAP regimens on the equine gut microbiota, log 2-fold changes (log2FC) were calculated based on OTU count changes on bacterial family level between sampling timepoints. Overall, the most prominent log2FC between t_0_ and t_1_ in the SSG were noticed for *Bacteroidaceae* (+5.16; p<0.05), *Enterobacteriaceae* (+3.99; p<0.05) and *Pseudomonadaceae* (+4.41; p<0.05). Among the most prominent log2FC between t_0_ and t_1_ in samples of the 5DG were *Bacteroidaceae* (+2.96; p<0.05) and *Pseudomonadaceae* (+3.33; p<0.05). In order to further investigate the relevance of the aforementioned observations and to better assess the influence of the colic syndrome complex as well as antibiotic treatment on the abundances of specific bacterial families in the GIT of horses throughout this study, further log2FC values were determined based on the abundances calculated for the CG.

#### Divergence among microbiota composition on family level considering a baseline defined by the CG samples

Since the PCA confirmed the eligibility of the CG samples with respect to an overall favourable equine gut microbiota structure and composition, we determined the median relative abundance for bacteria (family level) among the CG samples. To enhance identification of relevant changes associated with microbiome disturbance among samples belonging to both study groups, the median for each bacterial family was calculated using the t_0_ CG samples as a baseline. Then, log2FC for each sample/timepoint/study group/ were determined to pin-point variation that might have been overlooked by simply investigating mean abundances. In order to compensate for individual differences between the equine patients (i.e. age, diet, medical history, severity and duration of the acute colic episode, housing and social contacts), an interval comprising at least 85% of the SSG and 5DG samples was defined (**Figure 4**; log2FC +/- 2.5).

**Figure 4.**
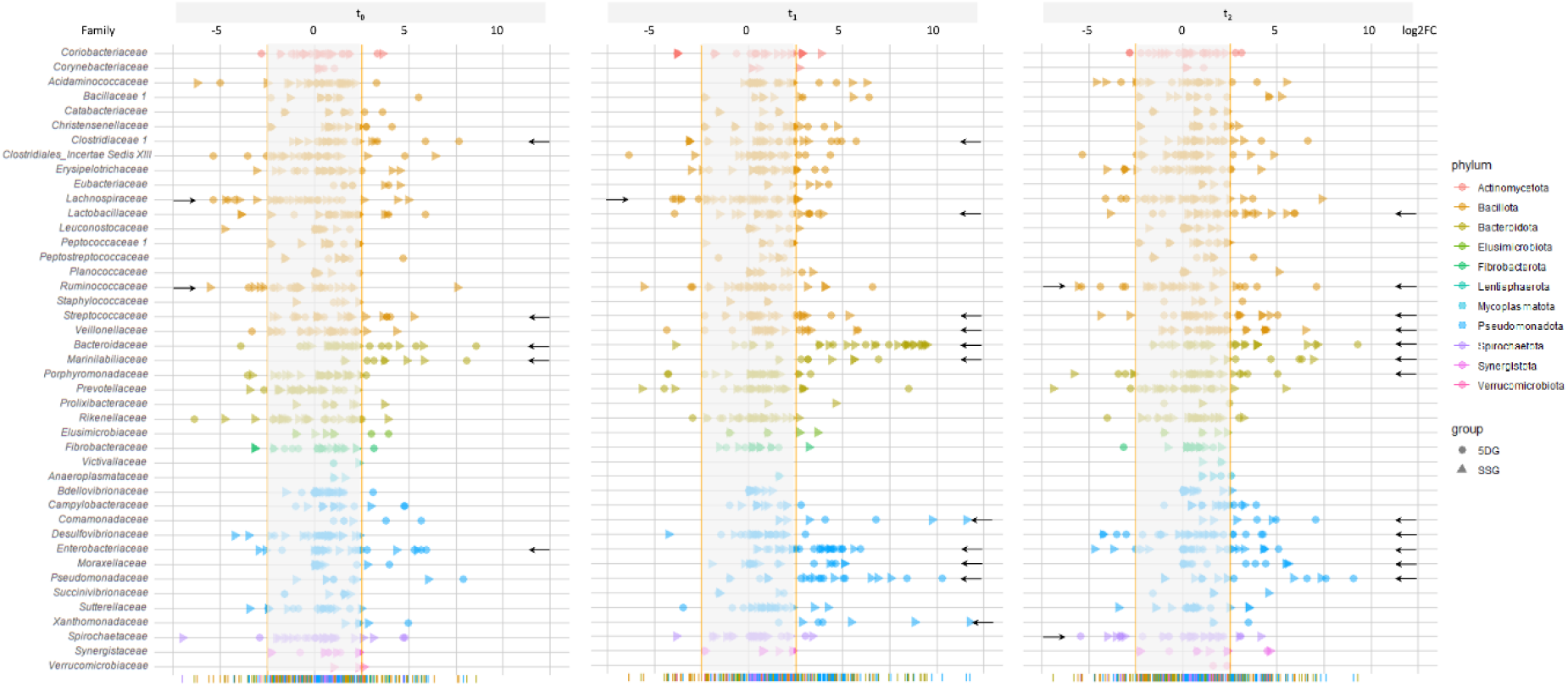
Changes of the fecal microbiome structure and composition during the study period. Illustration of log2FC abundance changes for bacterial families in each sample of both study groups with respect to a baseline calculated from the control group samples at t_0_. To acknowledge individual variation and to enhance the chances of overall trend detection, an interval comprising the majority of samples (85% at t_0_; log2FC of 2.5) was calculated, illustrated as the blurred area per timepoint. Individual values that were zero (i.e. no abundance difference compared to the control) were not shown for better clarification. Data points belonging to samples from the SSG are marked by a triangle and those from the 5DG by a dot, while deviation from the baseline within at least 5 samples are indicated by a black arrow.

To identify the most deviations with high potential relevancy, further in-depth explorative analysis was restricted to bacterial families associated with at least five samples (SSG & 5DG) that clustered beyond the 85% interval (**Figure 4**, black arrows indicate bacterial families fulfilling the restriction criteria).

Directly at hospital admission and before any antibiotics were administered (t_0_), six samples showed log2FC between 2.7 and 5.2 for *Streptococcaceae*, seven samples for *Bacteroidaceae* (log2FC between 2.9 and 8.6), seven samples for *Marinilabiliaceae* (log2FC between +2.8 and +8.1) and six samples for *Enterobacteriaceae* (log2FC between +2.8 and +5.9). Overall, lower values were noted for *Lachnospiraceae* (six samples, log2FC between -3.1 and -5.4) and *Ruminicoccaceae* (five samples, log2FC between -2.7 and -3.5) (**Figure 4**, t_0_; indicated by black arrows).

At t_1_, increasing log2FC fulfilling the above mentioned criteria were noticed for *Clostridiaceae*, *Lactobacillaceae*, *Streptococcaceae*, *Veillonellaceae*, *Marinilabiliaceae* and *Xanthomonadaceae*, but most prominently for *Bacteriodaceae* (22 samples, log2FC between +3.7 and +9.5), *Comamonadaceae* (five samples, log2FC between +3.2 and +11.6), *Enterobacteriaceae* (17 samples, log2FC between +2.7 and +5.9), *Moraxellaceae* (6 samples, log2FC between +3.5 and +5.1) and *Pseudomonadaceae* (15 samples, log2FC between +2.8 and +10.3) (**Figure 4**). *Lachnospiraceae*, on the other hand, yielded log2FC between -2.7 and -4.0 in five samples.

At t_2_, deviation from the baseline interval (log2FC ≥+/- 2.5) defined for bacterial families were recognized for *Bacteriodaceae* (eleven samples, log2FC between +2.8 and +9.3), *Enterobacteriaceae* (eight samples, log2FC between +2.6 and +5.1) and *Lactobacillaceae* (nine samples, log2FC between +2.8 and +5.9) followed by *Streptococcaceae*, *Veillonellaceae*, *Marinilabiliaceae*, *Comamonadaceae*, *Moraxellaceae* and *Pseudomonadaceae* (**Figure 4**). Overall, a shift back to the baseline was directly recognizable for most of the displayed bacterial families.

Taken together, OTU abundances of many bacterial families deviated considerably from those associated with samples of the CG (t_0_). In addition, overall deviation increased at t_1_ and decreased at t_2_, a result that is clearly in congruence with the SDI values and the trajectory pattern displayed by PCA (**Figure 3A-D**). Of note, clear differences between the study groups were not noticed in **Figure 4**. These results confirmed that although a common temporal trajectory pattern was recognizable, deviation of individuals contributed largely to the overall variances.

### iii) Hospitalization, surgery and administration of PAP is accompanied by a predominant converged trajectory pattern of *Escherichia* and *Bacteroides*

At t_1_, deviations of *Bacteroidaceae* and *Enterobacteriaceae* stood out considering the baseline defined by the CG samples (**Figure 4**). Although relative abundances of OTUs within an individual sample depended on each other, we noticed stunning trajectories of OTUs assigned to *Bacteroides* and *Escherichia*, the predominating genera of the above-mentioned families within our sample set **(Figure 5**).

**Figure 5.**
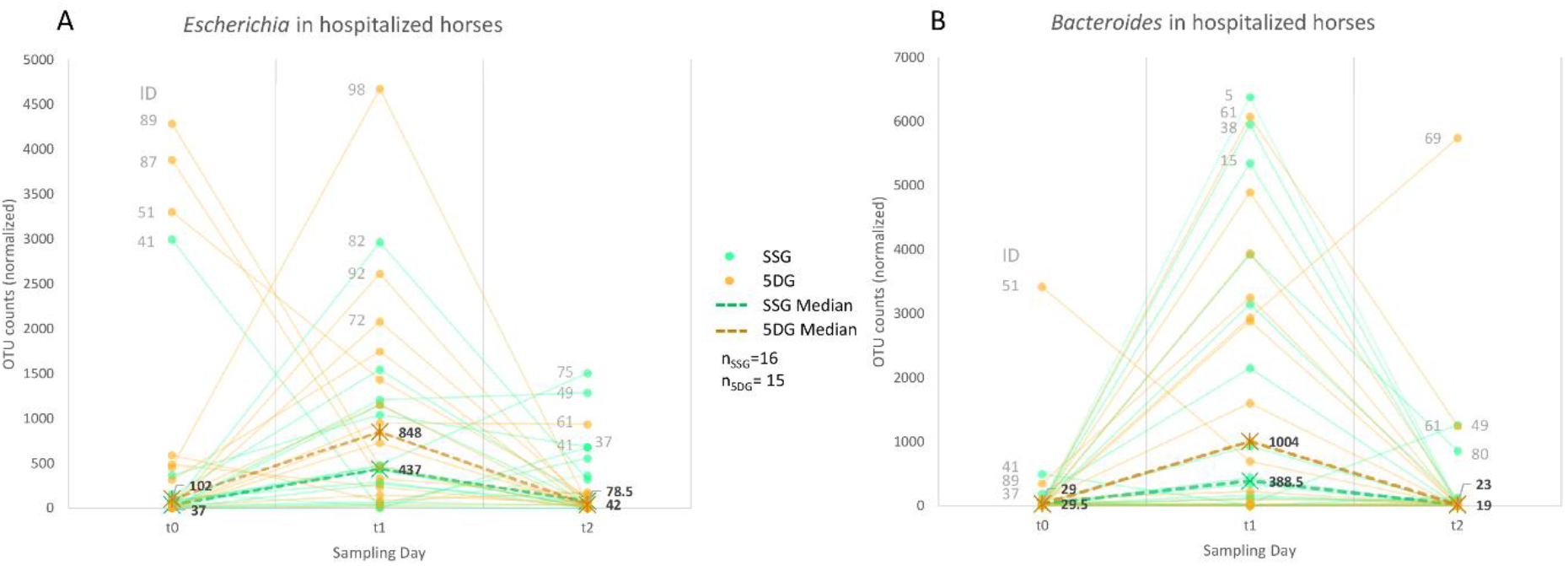
Abundance trajectories of OTUs assigned to the genera Escherichia and Bacteroides. Charts showing a trend line based on normalized OTU counts per sample/study group and timepoint. A dashed line indicates the median per group and sampling day.

Apart from four horses (89, 87, 51 and 41), most study participants showed limited OTU counts (median_SSG_= 37; median_5DG_= 102) classified as *Escherichia* (genus level) at t_0_, while the overall counts increased at t_1_ (**Figure 5A**), with a median OTU counts of 437 (SSG) and 848 (5DG) in samples of both groups, respectively (SSG vs. 5DG, t_0_: p=0.14, t_1_: p=0.086, t_2_: p=0.95). This difference was significant between the SSG and 5DG from t_1_ to t_2_ (differences in counts, SSG vs. 5DG, t_0_ to t_1_: p=1, t_1_ to t_2_: p=0.0019). Of note, both study groups showed a decline to almost similar counts assigned to *Escherichia* at t_2_ (median_SSG_= 79 OTU counts; median_5DG_= 42 OTU counts).

Interestingly, the hospitalized horses demonstrated a highly similar trajectory pattern for OTU counts assigned to the genus *Bacteroides*. Besides a single exception (horse 51, 5DG), both study groups had comparable counts for *Bacteroides* (median_SSG_ = 29.5 OTU counts, median_5DG_ = 29 OTU counts) at t_0_ (SSG vs. 5DG, t0: p=0.67, t1: p=0.45, t2: p=0.69). Changes in differences were not significant across time points (differences in counts, SSG vs. 5DG, t_0_ to t_1_: p=0.89, t_1_ to t_2_: p=0.92).

At t_1_, the OTU counts for *Bacteroides* doubled in abundance within the 5DG compared to the SSG (median_SSG_ = 389, median_5DG_ = 1004 OTU, p = 0.45). However, the *Bacteroides* count level decreased once more at the last day of sampling (t_2_) in both groups (median_SSG_= 19; median_5DG_= 23, p=0.69).

In summary, the abundance counts of both *Escherichia* and *Bacteroides* seemed to be associated with a more pronounced increase among samples representing the 5DG, i.e. horses that received the long-term PAP, compared to horses belonging to the SSG at t_1_. Correlation analysis revealed no obvious relationship between the two genera in the sample set investigated.

## 4 Discussion

In the present study, we explored temporal alterations of the gut microbiota composition and the extent of its perturbation among hospitalized horses subjected to colic surgery that received either a short-term (SSG) or a 5-day course (5DG) of P/G PAP. Of note, comparison of fecal microbiota profiles of horses belonging to different cohorts or even between distinct individuals requires immense caution, since the intestinal microbiome is easily affected by external factors, including - but not limited to - exercise (Górniak et al., 2021), transport, fasting (Schoster et al., 2016) and diet (Al Jassim, 2006;Mshelia et al., 2018). In addition, Antwis and colleagues recently examined how spatial and social interactions affected the gut microbiome composition of semi-wild Welsh Ponies, revealing that up to 52.6% of the observed variation is attributable to individual variation (Antwis et al., 2018).

At first, we determined the predominating microbial phyla in the sample set, revealing *Bacteroidota* (38%), *Bacillota* (33%) and *Verrucomicrobiota* (11%) as the top three main phyla at hospital admission (t_0_) and in the CG (**Figure 2**). This result is in line with previous reports defining the *Bacteroidota* and *Bacillota* as the major phyla of the core bacterial community in equines (Morrison et al., 2018;Edwards et al., 2020).

### Gut microbiota diversity of hospitalized horses receiving colic surgery and P/G PAP elicits a predominant trajectory pattern from hospital admission to discharge

As summarized previously, most of the current clinical studies regarding the effects of antibiotics on the gut microbiome have been cross-sectional, while interventional or longitudinal approaches and comparisons to treatment-naive but diseased control groups are often missing (Zimmermann et al., 2021). As a result, it is difficult to differentiate between disease-mediated, drug-related hospitalization effects (Zimmermann et al., 2021), an aspect that clearly is a limitation with respect to the discussion of putative drug-related effects in the current study. Although the isolated and exact impact of the P/G regimen on the gut microbiota of horses participating in this study remains unknown due to the fact that our study was not an animal trial but a real-world scenario, recent research on the effect of parental administration of P/G on the developing infant gut microbiome clearly revealed an impact on Shannon diversity and overall gut microbiota composition (Reyman et al., 2022). However, at hospital admission and before antibiotic treatment and hospitalization (t_0_), the mean alpha-diversity measured by SDIs revealed strong signals of an already compromised bacterial biodiversity in samples representing the SSG and the 5DG compared with those of the CG (**Figure 3D**), although only the latter difference was found to be significant. A loss of species diversity seems to be among the prominent characteristic of a disturbed gut microbiota (Ramirez et al., 2020). At hospital admission, a decrease in bacterial richness and diversity accompanied with a greater inter-individual variability was reported for horses admitted due to colic compared with horses presented for elective surgical procedures or even healthy horses (Stewart et al., 2018;Park et al., 2021). A similar trend was detected in the current study, demonstrating a considerable range of variation between microbiota profiles and microbiome perturbations among horses suffering from colic compared to horses free of abdominal pain or otherenteral disorders.

A previous study by Costa et al. examined the effects of intramuscular administration of procaine penicillin and ceftiofur sodium, and oral trimethoprim sulfadiazine on the fecal microbiome of healthy horses (Costa et al., 2015). Although the administration route and duration, the cohort under investigation and the combination of antibiotics (P/G) differed in the present study, general similarities should be mentioned. The strongest perturbation of the microbiome was recognized directly after the final course of antibiotics, including a strong effect on the microbial community membership (Costa et al., 2015).

The most prominent peak of microbiome disturbance is characterized by significant SDI decreases at t_1_ (i.e. three days after surgery/hospitalization) (**Figure 1**). The considerable inter-individual distances between the microbiota profiles of 5DG horses at t_0_ have diminished immensely at t_1_, indicating a similar and consistent effect of the long-term PAP regimen on the gut microbiota composition that is specific for the long-term P/G PAP.

This observation is in line with the results of a recent study investigating the effects of commonly used antibiotics on the gut microbiome of healthy human volunteers before and after treatment, where the authors revealed drug-specific trajectories through the PCA space over time (Anthony et al., 2022). In strict contrast to the PCA of the 5DG in the present study, data points of the SSG showed increased inter-individual differences through the PCA space at t_1_, indicating the lack of a common selective factor. Therefore, other more distinguishing factors associated with the distinct individual such as appetite, stress and/or pain, environmental bacteria or even the onset of GIT function and/or microbiome recovery (reviewed in (Kauter et al., 2019)) might be mirrored here.

At t_2_, both study groups showed clear signs pointing towards the onset of microbiome recovery with respect to a rise of both SDI_mean_, with most participants seeming to have already regained the baseline condition, since the difference between t_0_ and t_2_ lacked statistical differences (**Figure 3**). Interestingly, in a study of germ-free mice, the recovery of the gut microbiome after antibiotic treatment strongly depended on diet, community context and environmental reservoirs (Ng et al., 2019). The authors demonstrated that a reduction of environmental reservoirs impaired the process of microbiota recovery (Ng et al., 2019). This fact not only emphasizes the overall importance of the actual environmental bacteria in the immediate vicinity of hospitalized horses during microbiome recovery, but it also highlights the susceptibility of the equine gut microbiota to spatio-temporal local spreads of hospital-associated pathogens, explicitly including ESBL-EC, leading to worrisome carriage rates, as previously reported (Kauter et al., 2019).

Since studies on equine gut microbiomes are currently limited (Ang et al., 2022), it seems reasonable to assume that some general effects occur across different mammalian species: Exposure of gut microbiota to antibiotics or their still active metabolites reduces its diversity, while the absence of antibiotics leads to an altered state that is either transient or permanent (Costa et al., 2015;Ramirez et al., 2020;Arnold et al., 2021). Moreover, the effect might not be limited to a reduced microbial diversity, since a recent study on healthy human volunteers showed a worrisome increase of virulence- and resistance associated-factors immediately after antibiotic treatment (Palleja et al., 2018). Further studies on the equine gut metagenome (Ang et al., 2022) after exposure to different antibiotics are required to gain more insights with respect to spatial trends of virulence- and resistance gene occurrences and abundances.

### Alterations of the equine gut microbiota in the SSG and 5DG

At first, we analyzed the significant log2FC of the SSG and the 5DG. Samples of horses belonging to both study groups were associated with a significant abundance increase for *Bacteroidaceae*, *Enterobacteriaceae* and *Pseudomonadaceae* from t_0_ to t_1_. A previous study that examined changes in the equine fecal microbiome during hospitalization because of colic reported a similar increase for *Bacteroides* (Stewart et al., 2021). The increased abundances noticed for *Enterobacteriaceae* at t_1_ in both groups might enhance the risk of developing surgical site infections (SSI) for the equine patients, especially since multidrug-resistant Enterobacterales were often reported within horse clinics (Apostolakos et al., 2017;Walther et al., 2018;Shnaiderman-Torban et al., 2020), respectively in cases of SSI of horses subjected to laparotomy (Isgren et al., 2017;Dziubinski et al., 2020). Furthermore, significant changes were recognized regarding *Pseudomonadaceae*, a bacterial family that has been described as indicator of intestinal microbiome alterations in the human gut since an abundance increase seems to be associated with various gastrointestinal diseases (Alam et al., 2020;Chamorro et al., 2021).

Secondly, we explored the probable influence of the acute disease and other factors on the fecal microbiota. For this, we evaluated the abundance of different bacteria in fecal samples of both study groups compared to the control group at t_0_, revealing considerable deviation of OTUs belonging to 15 different bacterial families (**Figure 4**). Overall, a common temporal trajectory pattern was recognizable regarding *Bacteroidaceae*, *Enterobacteriaceae* and *Pseudomonadota*, with a strikingly increased proportion in almost all t_1_ samples and decreasing, but still above-baseline level, OTU counts at t_2_. In addition, we observed a reduced frequency of *Lachnospiraceae* for the horses diagnosed with colic, compared to the horses belonging to the CG, which is in line with recent findings (Stewart et al., 2019). Moreover, a reduction of *Lachnospiraceae* was previously described for horses suffering from enteral disorders (Costa et al., 2012;Weese et al., 2015).

Although differences between the study groups were limited, we observed interesting differences with respect to the effect of the distinct P/G PAPs on the microbiota. Horses that received the 5-day course of PAP showed the highest loss of inter-individual diversity (**Figure 3B**, t_1_) which is clearly accompanied by the most prominent expansion of *Escherichia* (**Figure 5**). Further studies including metagenomic data are needed to reveal the particular dependences here.

### Gut microbiome perturbances are associated with increased *Escherichia* and *Bacteroides* abundances

Since penicillin and gentamicin were administered parenterally (Stöckle et al., 2021), the effect on the gut microbiome in general was expected to be lower than in cases when oral antibiotics where administered (Zhang et al., 2013). In the present study, however, the combined influences of (intestinal) illness, hospitalization, surgery and administration of P/G PAP showed a converged relative abundance trajectory of the genera *Escherichia* and *Bacteroides* over time, as demonstrated in the results (**Figure 5**). A recent review on the effects of distinct antibiotic classes on the GIT microbiome in humans highlighted an increase of *Enterobacteriaceae* and *Bacteroidaceae* after treatment with β-lactams in general (Patangia et al., 2022), which seems strikingly in line with the results presented here.

The obligatory anaerobic genus *Bacteroides* exclusively growing in the GIT of mammals is a major research focus of gut microbiology (reviewed (Wexler and Goodman, 2017)). Since intrinsic resistance is reported for *Bacteroides* spp. (Pumbwe et al., 2006;CLSI, 2020) at least an impact of a treatment-associated selective advantage can be assumed when considering the relative abundance increase among most of the fecal samples of both study group participants at t_1_ (**Figure 5**), in particular for the 5DG.

Although many *Bacteroides* species play a crucial role in degrading polysaccharides of a plant-based diet (Pereira et al., 2021;Cheng et al., 2022), the specific importance and role of distinct *Bacteroides* species in hindgut fermentation has not been investigated yet. More research on the subject including metabolomic profiles gained from metagenome sequencing projects is clearly required to shed more light on this subject.

Apart from outliers (four horses, **Figure 5**), most of the horses participating in the present study revealed a low relative abundance of *Escherichia* at hospital admission. This observation is in line with the previously reported increase of ESBL-EC carriage rates among patients of both study groups over time, i.e. from hospital admission to discharge (**Supplementary Table 1**), that indicated local spread of, for instance, distinct ESBL-EC clonal lineages, as reported previously (Kauter et al., 2021). We further speculate that during P/G PAP, ESBL-EC had a selective advantage in the GIT of horses and therefore proliferate, resulting not only in increased carriage rates (Kauter et al., 2021), but also in unavoidable environmental contamination via feces-contaminated litter. Since the immediate environment is among the main sources of GIT-associated bacteria during microbiome recovery (Ng et al., 2019), environmental sources seem to play an overwhelming role with respect to ESBL-EC carriage rates. Since the specific needs and surroundings of hospitalized horses (bedding, boxes, floor surfaces, reviewed in (Gehlen et al., 2020) differ immensely from those of small animal or even human patients, the environment of equine clinics seems to play an important role in ESBL-EC accumulation and spread, including strains known for their pathogenic potential in humans and animals, i.e. isolates belonging to sequence type (ST)10 and ST410 (Kauter et al., 2021).

Taken together, the spread of *E. coli*, especially ESBL-EC in horse clinics, seems to be promoted by i) the selective advantage of these bacteria towards β-lactam antibiotics and ii) the fact that the fecal microbiota structure is re-modelled by other factors occurring during the course of hospitalization, such as ingestion by food, contact to environmental sources or transmission via healthcare workers.

## Conclusion

In the present study, we investigated the influence of two different PAP regimens (SSG vs. 5DG) in horses diagnosed with colic syndrome that were subjected to surgery with regard to changes of their gut microbiota composition. Colic surgery and PAP drive the equine gut microbiome towards dysbiosis and reduced biodiversity that is accompanied by a 10-fold increase of samples positive for ESBL-producing Enterobacterales (Kauter et al., 2021) and an abundance increase for Enterobacteriaceae. Further studies are needed to reveal the most important local sources of the resistant bacteria (i.e. environment, food, contacts) and factors promoting the inclusion of ESBL-producing Enterobacterales in the equine gut microbiota.

## Supporting information

Supplemental Figure 1_Barplot_phyla_all

## Availability of data and materials

The established workflow was implemented in Python and is freely available under GPLv3 license (https://github.com/SiWolf/Meta16s/). The repository includes further downstream analysis and visualization scripts in R, as well as an associated Conda environment for reproducibility. Raw 16S rRNA gene sequences were submitted to NCBI and are stored within BioProject PRJNA906950.

## Authors’ contributions

BW, AL-B, LHW and HG designed the project. AK, SDS, HG, and BW conceived and designed the experiments. AK, DK, SDS, JB and AL-B performed laboratory analysis. CB and SF sequenced the samples. AK, SAW, AL-B, BW and TS analyzed the data. AK, SAW, and BW wrote the first draft. AM, NE, RK and SG helped to draft the manuscript and contributed to the discussion. All authors have read and approved the final draft of the manuscript.

## Ethics

According to the German regulation authorities for research with animal subjects, the comparison of two PAP regimens does not require explicit approval (Landesamt für Gesundheit und Soziales, Berlin, 18.04.2017). Written owner’s consent regarding the involvement of their horses in the study was obtained directly during the hospital admission process (Stöckle et al., 2021).

## Acknowledgements

We thank our colleagues from the Advanced Light and Electron Microscopy (ZBS 4) department of the Robert Koch Institute for their individual contribution and support.

## Abbreviations

5DG: 5-day course of peri-operative antibiotic prophylaxis group
Bp: base pair
CG: control group
EC: *Escherichia coli*
ESBL: extended-spectrum β-Lactamase
G: gentamicin
GIT: gastrointestinal tract
i.e.: id est
Log2FC: log_2_ fold change
MDR: multi-drug resistance
OTU: Operational Taxonomic Units
P: penicillin
PAP: perioperative antibiotic prophylaxis
PCA: principal component analysis
SDI: Shannon diversity index
SSG: single-shot of peri-operative antibiotic prophylaxis group

