## Supplementary figures and images for "Antibiotic prophylaxis and hospitalization of horses subjected to median laparotomy: gut microbiota trajectories and abundance increase of *Escherichia*"

### Supplemental Figure 1_Barplot_phyla_all

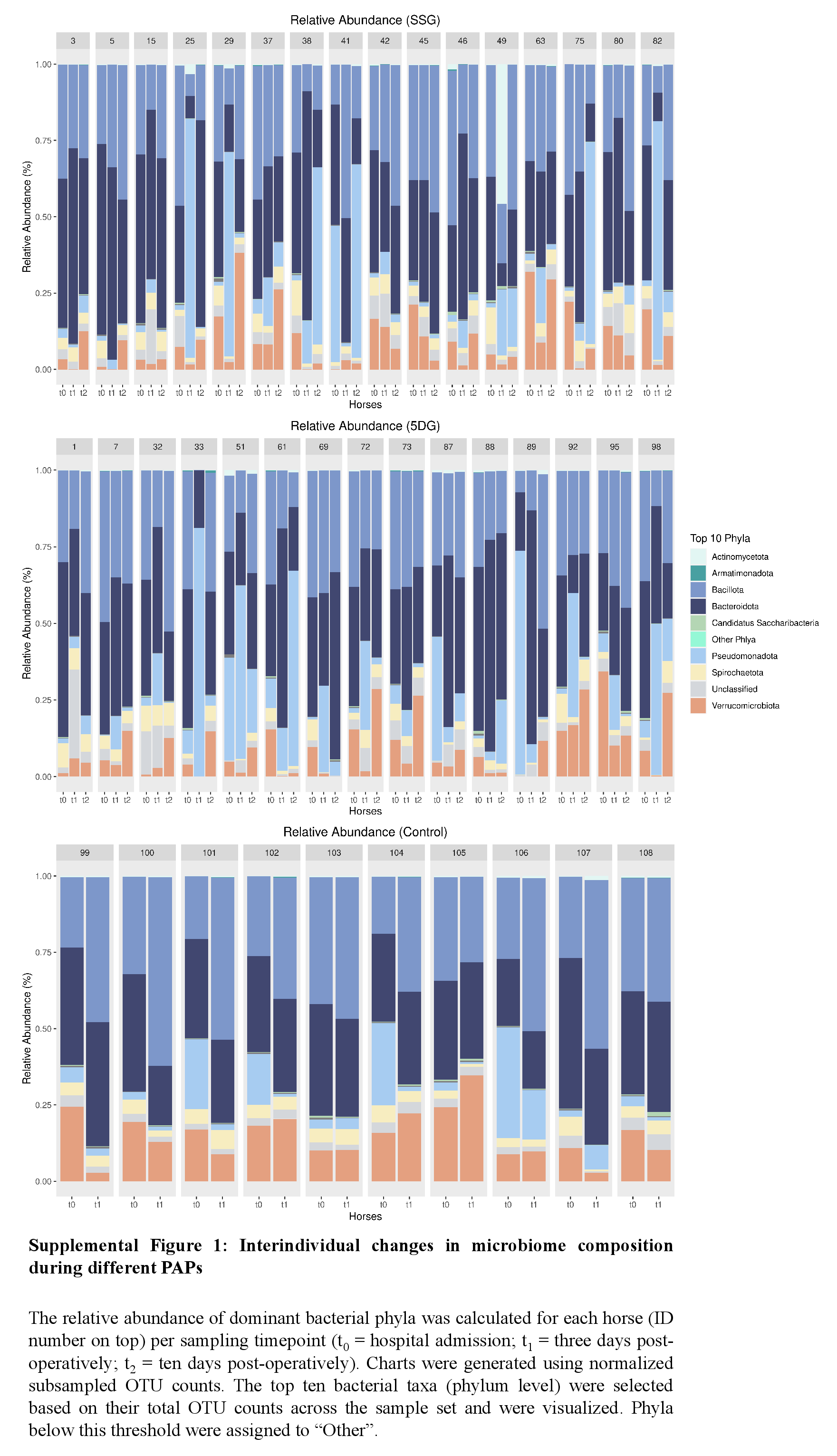
